# Potential impact of sexual transmission of Ebola virus on the epidemic in West Africa

**DOI:** 10.1101/031880

**Authors:** Jessica L. Abbate, Carmen Lia Murall, Heinz Richner, Christian L. Althaus

## Abstract

Sexual transmission of Ebola virus disease (EVD) 6 months after onset of symptoms has been recently documented, and Ebola virus RNA has been detected in semen of survivors up to 9 months after onset of symptoms. As countries affected by the 2013–2015 epidemic in West Africa, by far the largest to date, are declared free of Ebola virus disease (EVD), it remains unclear what threat is posed by rare sexual transmission events that could arise from survivors. We devised a novel mathematical model that includes sexual transmission from convalescent survivors: a SEICR (susceptible-exposed-infectious-convalescent-recovered) transmission model. We fitted the model to weekly incidence of EVD cases from the 2014–2015 epidemic in Sierra Leone. Sensitivity analyses and Monte Carlo simulations showed that a 0.1% per sex act transmission probability and a 3-month convalescent period (the two key unknown parameters of sexual transmission) create very few additional cases, but would extend the epidemic by 83 days [95% CI: 68–98 days] (p < 0.0001) on average. Strikingly, a 6-month convalescent period extended the average epidemic by 540 days (95% CI: 508–572 days), doubling the current length, despite an insignificant rise in the number of new cases generated. Our results show that current recommendations for abstinence and condom use should reduce the number of sporadic sexual transmission events, but will not reduce the length of time the public health community must stay vigilant. While the number of infectious survivors is expected to greatly decline over the coming months, our results show that transmission events may still be expected for quite some time as each event results in a new potential cluster of non-sexual transmission. Precise measurement of the convalescent period is thus important for planning ongoing surveillance efforts.

## Introduction

Recent reports suggesting the potential for sexual transmission of Ebola virus from convalescent survivors have raised a number of important questions regarding its impact on the final phase of the epidemic in West Africa [1,2]. Even once the worst hit countries of Guinea, Liberia, and Sierra Leone are declared free of Ebola virus disease (EVD), rare cases may still arise from the large number of remaining survivors. Importantly, sexual transmission is dependent on the frequency of infections rather than the density of available hosts, allowing chains of transmission to persist at low susceptible densities where non-sexual transmission would typically fail to occur [3]. Perhaps the most crucial element for bringing the epidemic to an end is maintaining vigilance in the community and preventing – or quickly responding to – new chains of transmission. Thus, it is important to investigate the potential impact of convalescent sexual transmission on the transmission dynamics in general, and on the tail of the epidemic in particular, to understand how long that vigilance might remain critical.

Follow-up studies on survivors of the 1995 outbreak in the Democratic Republic of Congo [4] and the 2000 [5] and 2007 [6] outbreaks in Uganda have raised awareness of what is now being termed “post-Ebola syndrome” (post-Ebola sequelae) – debilitating illnesses from myalgia to uveitis – which can persist for at least 21 months after the onset of symptoms. Though the virus is no longer detected in the blood after acute EVD symptoms disappear, active (replicating) virus has been documented in ocular fluid, rectal fluids, vaginal fluids, and semen [1,4,7,8]. Transmission to sexual partners was never confirmed in earlier outbreaks, but was suspected to have occurred in at least one instance [4]. Similarly, cases of sexual transmission of other hemorrhagic fever infections, notably by the closely related Marburg virus, have been suspected in the past [9,10]. Studies from the West African outbreak, showing viremia in semen 4–6 months after onset of symptoms in 65% of men tested (7–9 months in 26%) [1] and presenting molecular evidence of sexual transmission from a survivor 179 days after onset of symptoms [2], suggest that sexual transmission from convalescent men can and does occur.

Sexual transmission of Ebola virus from convalescent survivors is likely a rare event, but researchers have warned that it should be considered in epidemiological models that are used to predict the trajectory of an outbreak [11]. To this end, we devised a novel mathematical model for EVD transmission: SEICR (susceptible-exposed-infectious-convalescent-recovered), which includes a component for convalescent sexual transmission from convalescent survivors who maintain active Ebola virus replication.. We illustrated the model by fitting it to weekly EVD incidence from Sierra Leone, the largest population of recovering survivors from the current West Africa epidemic. We performed sensitivity analysis to understand the influence of key unknown parameters, such as the duration of the convalescent period and the transmission probability per sexual contact. We also performed Monte Carlo simulations to explore the impact of sexual transmission on the epidemic tail in Sierra Leone.

## Methods

***Transmission model***. We extended a SEIR (susceptible-exposed-infected-recovered) modeling framework, which has been extensively used to describe EVD transmission [12–14], by adding a component for sexual transmission from convalescent survivors who maintain active Ebola virus replication (Fig. 1). The resulting SEICR model has five states: susceptible, *S*, exposed, *E*, symptomatic and infectious, *I*, convalescent, *C*, fully recovered and immune, *R*, and dead, *D*. The model is represented by the following set of ordinary differential equations (ODEs):

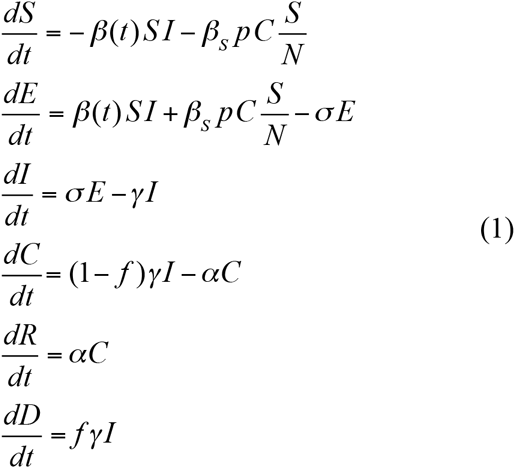

where *N = S + E + I + C + R* denotes the total population size. We assumed the nonsexual transmission rate, *β(t)*, to be initially constant (*β*_0_) before it starts to decay exponentially due to the effect of control interventions and behavior change after time *τ*: *β(t) = β*_1_ + (*β*_0_ - *β*_1_)e^−k(t−τ)^ [12]. The sexual transmission parameter, *β*_s_, can be described as the product of two parameters (*β*_s_ = *ηq*) that we will consider separately: *η* is the per sex act transmission probability of Ebola virus from convalescent men, and *q* is the daily rate at which they engage in sexual intercourse. The number of convalescent individuals are scaled by *p*, which is the proportion of convalescent survivors who are sexually active men. 1/*σ* and 1/*γ* represent the average durations of incubation and symptomatic infection, respectively. *f* is the case fatality rate. The average duration after which convalescent patients recover completely and shed no further replicating Ebola virus from their body is given by 1/α. We assumed that sexual transmission is frequency-dependent [3,15,16], i.e., the probability that the sexual partner of a convalescent man is susceptible is given by *S/N*.

**Figure 1.**
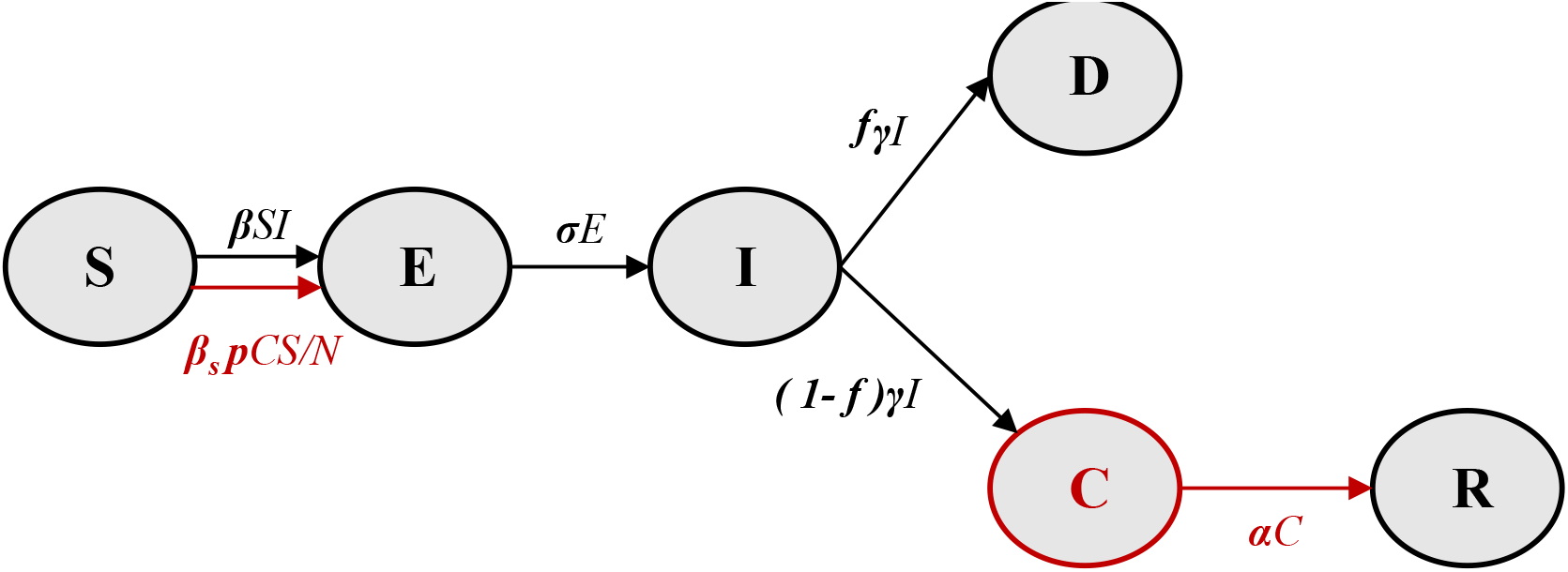
Schematic illustration of EVD transmission model including sexual transmission from convalescent patients. The elements in black form the base model without sexual transmission [12–14]. The red elements (convalescent individuals and additional transmission term) were added to account for sexual transmission.

The basic reproductive number, *R*_0_, for the SEICR model can be calculated using the next-generation matrix method [17,18] and is given by

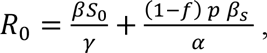

where *S*_0_ is the initial number of susceptible individuals (see *Supplementary Material* Appendix S1). When *α* goes to infinity or either *β*_s_ = 0 or *p* = 0, the equation reduces to the basic reproductive number in absence of sexual transmission: 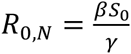. Thus, the second term represents the contribution of sexual transmission from convalescent patients to the overall 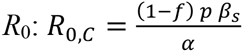.

***Model parameters***. We fitted the deterministic EVD transmission model to weekly incidence of confirmed and probable cases in Sierra Leone as reported in the WHO patient database [19] *(Supplementary Material* Fig. S1). The data set was extended with weekly incidence from the situation report for the most recent weeks when no data was available in the patient database. In order to account for variability in the accuracy of reporting, we assumed that the number of reported cases follows a negative binomial distribution with mean as predicted by the model and dispersion parameter *ϕ* [20]. We derived maximum likelihood estimates (MLE) of the following model parameters [14,21]: the baseline transmission rate *β*_0_, the time *τ* at which transmission starts to drop, the rate *k* at which transmission decays, and the dispersion parameter *ϕ*. For the fitting procedure, we assumed that there were no sexual transmission events, i.e., we set *β*_s_ to zero. The basic reproductive number in absence of sexual transmission is *R*_0,*N*_ = *β*_0_*N*_0_/*γ*, and the reproductive number in presence of partially effective control interventions is *R*_1,*N*_ = *β*_1_*N*_0_/*γ*, with *N*_0_ being the population size of Sierra Leone. We explored value ranges for sexual transmission parameters (Table 1) based on information from the current epidemic [22] and studies of human immunodeficiency virus [23,24]. Remaining parameters were based on published values from the literature (Table 1).

**Table 1:** Model parameters describing EVD transmission in Sierra Leone. The indicated parameter ranges are used for the sensitivity analysis (partial rank correlation coefficients, PRCC) only.

| Parameter |  | Value (Range) | Comments and References |
| --- | --- | --- | --- |
| Basic reproductive number without sexual transmission | $R_{0,N}$ | 2.13<br>(1.26 - 2.53) | Estimated - Range explores estimates from [13,25] |
| Basic reproductive number in presence of control interventions | $R_{1,N}$ | 0.67 | Estimated |
| Date of onset of symptoms in index case |  | April 23, 2014 | [13,25,26] |
| Rate at which transmission rate decays | $k$ | $0.011 \text{ d}^{-1}$ | Estimated |
| Time at which transmission rate starts to decay | $\tau$ | 51.0 d | Estimated |
| Incubation period | $1/\sigma$ | 11.4 d | [22] |
| Infectious period | $1/\gamma$ | 3.9 d | Together with the incubation period results in a generation time of 15.3 d [22] |
| Dispersion parameter | $\phi$ | 0.050 | Estimated |
| Initial population size | $N_0$ | $6.316 \times 10^6$ | Based on 2014 estimate [27] |
| Sexual transmission probability (per coital act) | $\eta$ | 0.001<br>(0.0005 – 0.002) | Roughly based on sexual transmission probability of HIV per coital act from infected men [24] |
| Case fatality rate | $f$ | 0.69 | [22] |
| Frequency of sex acts | $q$ | $0.272 \text{ d}^{-1}$<br>(0.136 – 0.544) | 8.27 coital acts per month [23] |
| Proportion of convalescent survivors who are infectious and sexually active | $p$ | 0.347<br>(0.1725-0.694) | Of 47.4% male survivors, 73.1% are aged 15-45 [22] |
| Rate at which convalescent survivors recover completely | $\alpha$ | $1/87.35 \text{ d}^{-1}$<br>($1/174.7 - 1/43.7$) | 3 months after onset of symptoms [4,28] and assuming an infectious period of 3.9 days |

***Deterministic model and sensitivity analysis***. We solved the system of ODEs numerically using the function *ode* from the *deSolve* package in the R software environment for statistical computing [29]. We compared the following response variables of the model: the epidemic peak number of exposed, *E*, acute, *I*, and convalescents, *C*, cases; the cumulative number of EVD cases, deaths, and recoveries; the date at the epidemic peak; the daily and cumulative incidence of sexual transmission; and the date at which the last symptomatic case either died or entered into convalescence (“day of last case”). We defined the day of last case as the time when the number of symptomatic and infectious individuals, I, dropped below 0.5. We considered the following parameters for the sensitivity analysis: the per sex act transmission probability of Ebola virus from convalescent men (*η*), the proportion of convalescent survivors who are sexually active men (*p*), the rate at which they engage in sexual intercourse (*q*), and the rate at which convalescent patients recover completely and shed no further replicating Ebola virus from their body (*α*). The sensitivity of the response variables to changes in *R_0_* was explored simultaneously as a comparison. We generated 1000 parameter combinations from the uniform ranges, log-transformed [0.5x – 2x] for the parameter values for *η, p, q*, and *α*, given in Table 1 via Latin hypercube sampling using the Huntington and Lyrintzis correlation correction method (function *Ihs* from R package *pse*) [30]. We then calculated partial rank correlation coefficients (PRCCs) using 50 bootstrap replicates [31].

***Monte Carlo simulations***. We performed stochastic simulations of the SEICR model with and without sexual transmission using Gillespie’s algorithm [32]. We specifically investigated the following response variables from the simulations: the cumulative number of EVD cases, the size and date of the epidemic’s peak incidence (daily number of new symptomatic infections), and the date of last case (last day that symptomatic infections, *I*, fell below 1). Summary statistics were based on the results of 1000 simulation runs for each transmission scenario. We calculated the average of the peak and total cumulative number of EVD cases by including all simulations runs, i.e., also the simulations that rapidly go extinct. In contrast, the average dates of the epidemic peak and last case were based on the simulated epidemic trajectories over which 50 or more cases were accumulated.

## Results

***Contribution of sexual transmission to overall R*_0_**. Assuming a conservative baseline scenario (*η* = 0.001 and 1/*α* = 3 months; Table 1), the reproductive number of a convalescent infection, *R*_0,*c*_, is 0.0024. This corresponds to only 0.12% of the overall *R_0_* of 2.0224. Increasing the convalescent period from 3 to 6 months, the contribution of *R*_0,*c*_ (0.0051) to the overall *R*_0_ rises to 0.25%. The equation for *R*_0,*c*_ (see *Methods*) illustrates that doubling the per sex act transmission probability has the same impact as doubling the convalescent period. It is important to note that the relative contribution of sexual transmission to the overall reproductive number rises as the effective reproductive number drops during the epidemic due to the effects of control interventions and decreasing density of susceptible hosts (see *Supplementary Material* Fig. S2).

***Effect of sexual transmission parameters on epidemic dynamics***. The two key unknown parameters of sexual transmission are the per sex act transmission probability, *η*, and the rate at which convalescent survivors fully recover, *α*. Both parameters were found to have very small effects on the peak number of infected or exposed patients (Fig. 2A; Fig. 3A; Figs. 4A, 4B, 4C; Fig. S2; Fig. S3). The duration of the convalescent period has a large impact on the peak number of convalescent individuals, while *η* does not (compare Fig. 2A and Fig. 3A). The total number of recovered individuals is reached more slowly the longer the convalescent period (Fig. 2B), which is not an effect caused by *η* (Fig. 3b). While the convalescent periods (1/*α* = [3 - 9 months]) and the values of *η*(*η* = [0.0005 - 0.002]) we explored create very few extra cases (Fig. 2B and Fig. 3B), sensitivity analyses revealed that a higher per sex act transmission probability, *η*, a higher proportion of sexually active convalescent individuals, *p*, or a higher frequency of sex acts, *q*, have larger impacts on the total number of cases than would proportional increases in the convalescent period (see *Supplementary Material*, Fig. S3A). Sensitivity analyses also revealed that these sexual transmission parameters could produce a small delay in the epidemic peak, more so than would changes in the convalescent period (see *Supplementary Material*, Fig. S3B).

**Figure 2.**
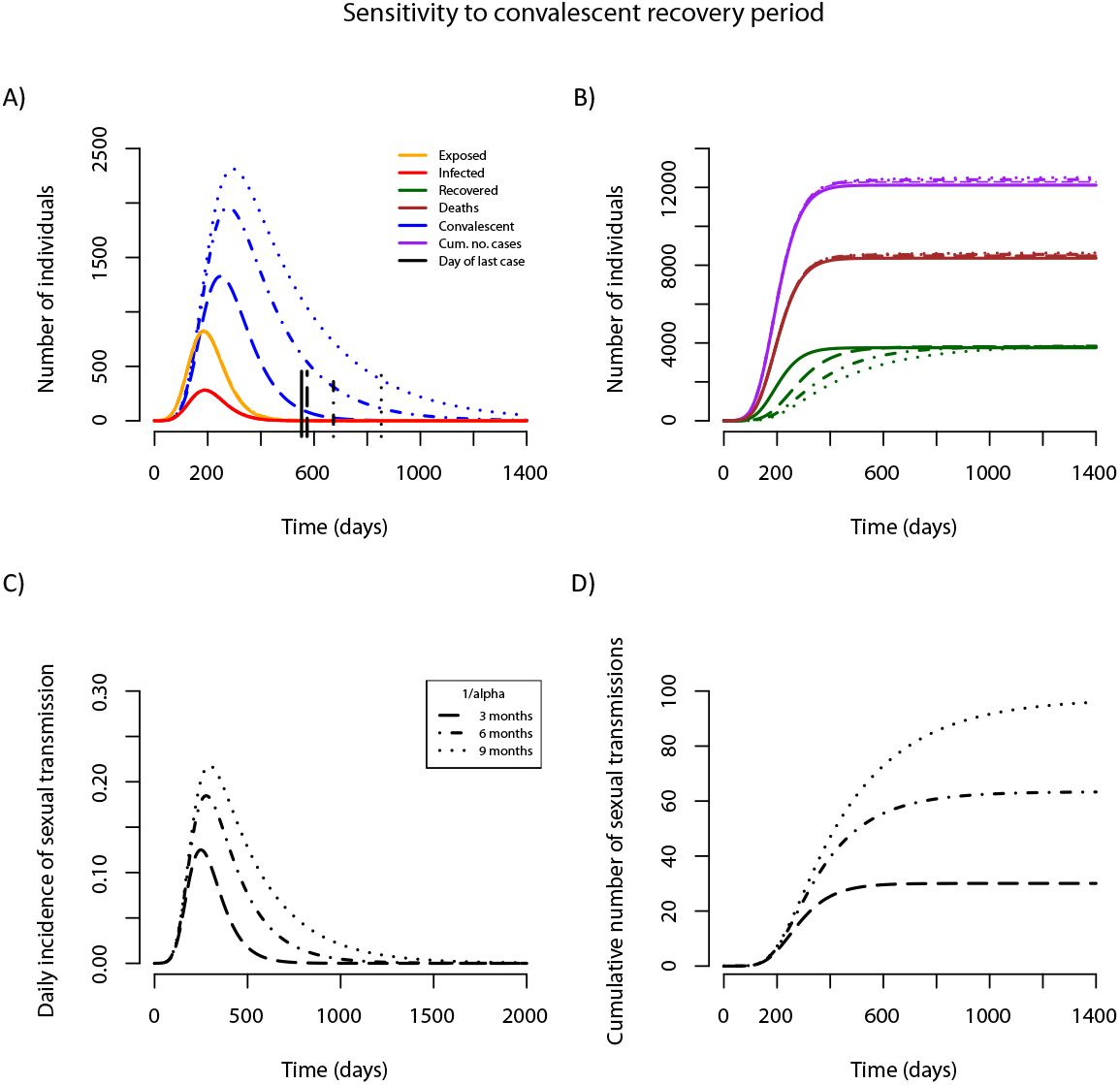
Effect of convalescent period on EVD epidemics. The average duration of the convalescent period (1/α) is varied between 3 and 9 months. (A, B): Epidemic trajectories in presence (broken lines) and absence of sexual transmission (solid lines). Vertical lines mark the day of last symptomatic case. (C) Daily incidence of sexual transmission. (D) Cumulative number of sexual transmission events. Note that the vertical axes vary across panels.

**Figure 3.**
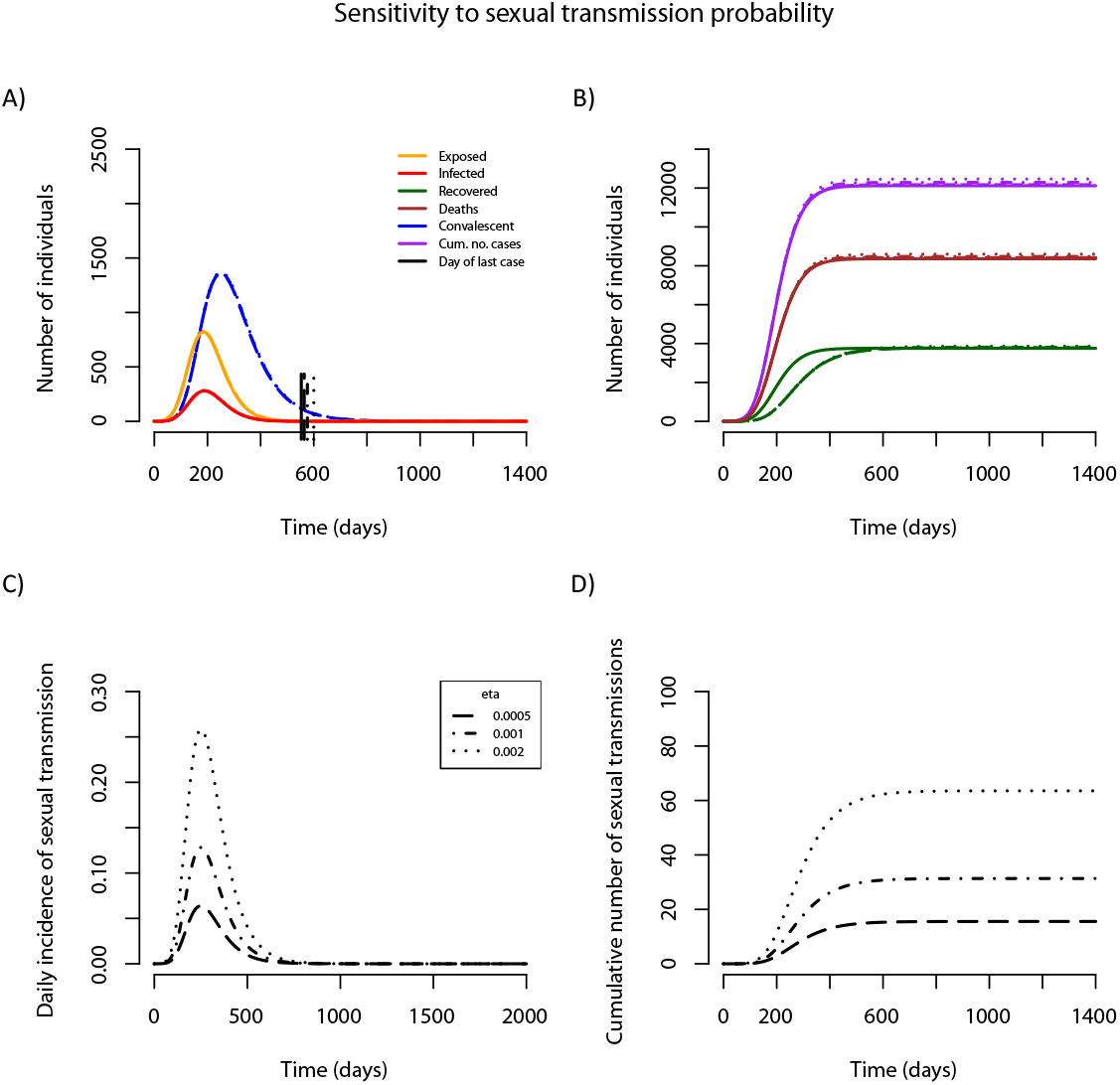
Effect of per sex act transmission probability on EVD epidemics. The per sex act transmission probability (*η*) is varied between 0.05% and 0.2%. (A, B): Epidemic trajectories in presence (broken lines) and absence of sexual transmission (solid lines). Vertical lines mark the day of last symptomatic case. (C) Daily incidence of sexual transmission. (D) Cumulative number of sexual transmission events. Note that the vertical axes vary across panels.

The number of sexual transmission events expected from the baseline scenario (*η* = 0.001 and 1/*α* = 3 months) is 31.2, the majority of which will occur around the peak of the epidemic (Fig. 2C and Fig. 3D) and thus likely go undetected. Doubling either *η* or 1/*α* results in nearly equal increases in the incidence and cumulative number of sexual transmission events (Fig. 2C, 2D and Fig. 3C, 3D), with either leading to roughly double the number of sexually transmitted cases over the course of the whole epidemic (> 60 cases). It should be noted that the total number of cases increases more than by simply the number of sexual transmission events, because each sexual transmission event results in a new potential cluster of non-sexual transmission. The day of last case is affected more by the convalescent period than the per sex act transmission probability (represented by vertical lines in Fig. 2A and Fig. 3A), a result confirmed by the sensitivity analysis (see *Supplementary Material*, Fig. S3A). The tail of the epidemic will depend on a small number of events that are likely to be affected by stochastic processes, thus we used Monte Carlo simulations to explore this behaviour.

***Impact of sexual transmission on the epidemic tail***. We performed stochastic simulations of the EVD transmission model to investigate the epidemic dynamics when the number of new cases becomes small, i.e, during the tail of the epidemic. Comparing model simulations while assuming a convalescent period of 3 months to those without sexual transmission confirmed the deterministic results that sexual transmission from convalescent survivors does not lead to a significant increase in the cumulative number of infected cases (non-STI: 11,092 +/- 627 cases; STI: 10,944 +/- 642 cases; Wilcox rank sum test: W = 491990, p = 0.53), nor the size (non-STI: 77 +/- 4.1 new cases per day; STI: 75 +/- 4.2 new cases per day; W = 493710, p = 0.62) or timing (non-STI: day 187 +/- 0.9; STI: day 187 +/- 0.9; t = 0.19, df = 1017.4, p = 0.85) of the epidemic peak incidence (Fig. 4A, 4B, 4C). This conservative period of potential sexual transmission, which has recently been shown to extend well beyond 3 months in at least 65% of patients [1], lengthened the average date on which the last active case could be detected by nearly three months (non-STI: 548 +/- 4.0 days; STI: 630 +/- 6.6 days; difference: 83 days [95% CI: 68–98 days], *t* = -10.8, df = 867.97, p < 0.0001; Fig. 4D, 4E, 4G, 4H). The width of the tail (s.d. = 151 days) was such that 23.4% of the 529 simulated epidemics that accumulated at least 50 cases still experienced symptomatic individuals 730 days (two years) after the start of the epidemic (Fig. 4H). Strikingly, when the convalescent period was extended from 3 months to 6 months, the projected length of the epidemic increased to a mean of 1088 days (+/- 15.5), with 84.0% of the 538 sustained epidemics taking over two years to end (Fig. 4F, 4I). However, the average number of new cases produced remained small (11,869 +/- 663 days; W = 482790, p = 0.18). Importantly, there is greater variance in the tail of the epidemic when sexual transmission is considered, and this uncertainty grows with the length of the convalescent period (Fig. 4G, 4H, 4I).

**Figure 4.**
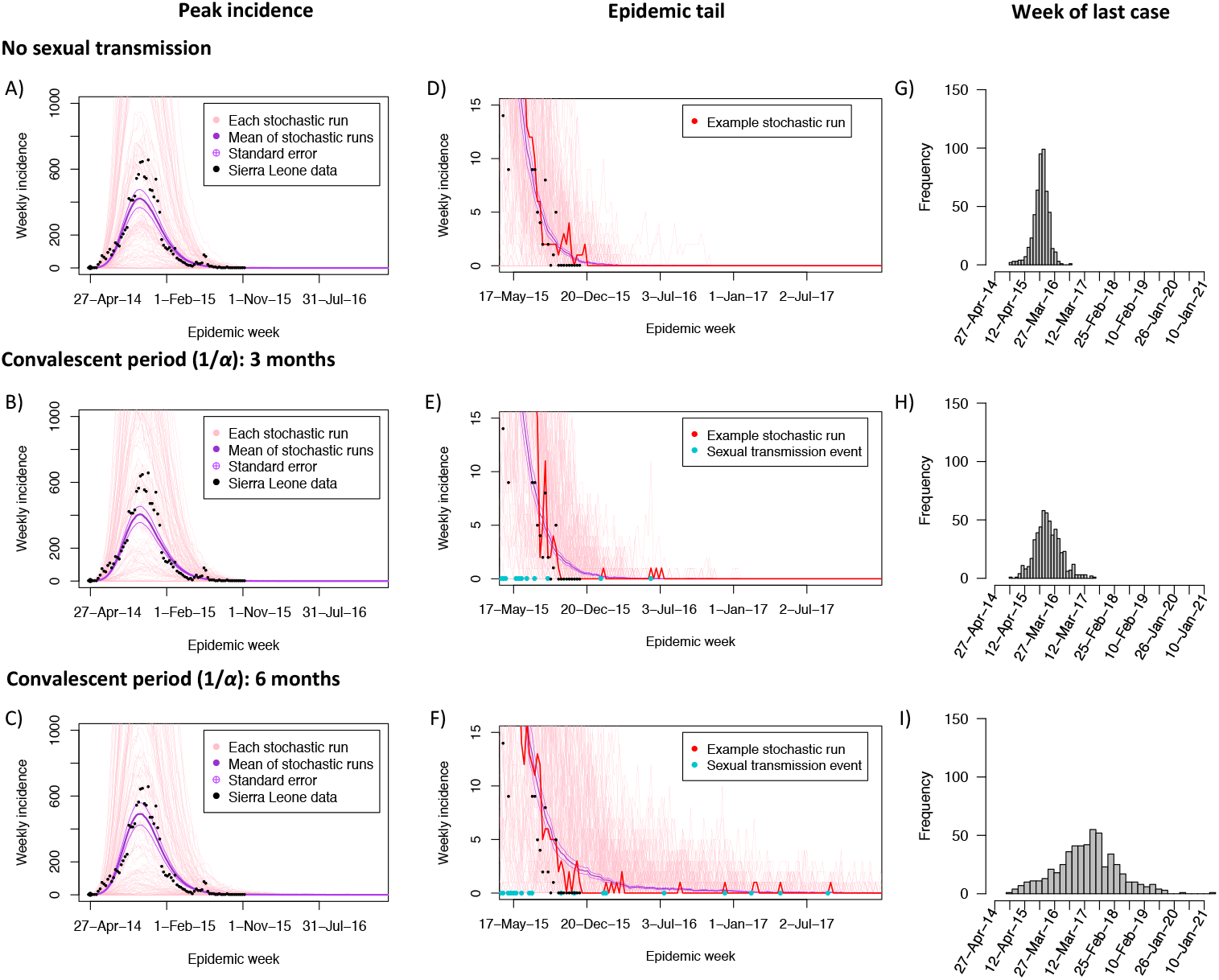
Impact of convalescent period on the tail of the EVD epidemic. Monte Carlo simulations of the weekly incidence of new cases (A, B, C) and the sporadic occurrence of sexual transmission events at the epidemic tail (D, E, F), assuming no sexual transmission (A, D), and sexual transmission with a 3 (B, E) and 6 months (C, F) convalescent period. Light red lines show the result of 200 simulated trajectories, with corresponding mean (thick purple line) and standard errors (thin purple lines). Black dots denote incident cases in Sierra Leone as reported by the WHO [19]. The dark red lines in (D), (E), and (F) highlight a single representative trajectory, and the blue dots along the horizontal axis indicate a sexual transmission event. Histograms of the day of the last EVD case from 1,000 simulated epidemics in absence of sexual transmission (G) and for a convalescent period of 3 (H) and 6 (I) months.

## Discussion

Our study shows that the length of the convalescent period will determine whether or not sexual transmission of Ebola virus from recovering patients will have a profound effect on the length of time it will take for the epidemic to completely fade. For Sierra Leone, we found that an average convalescent period of 3 months, and a per sex act transmission probability of 0.1%, could extend the EVD epidemic in Sierra Leone by an average of 83 days (95% CI: 68–98 days). Such a scenario would be consistent with the occurrence of a small number of sexual transmission events during the end-phase of the epidemic. However, assuming an average convalescent period of 6 months led to simulated epidemics whose tails were much more variable, and much longer, despite a lack of significant increase in the total number of cases. So far, the reported cases of sexual transmission of EVD remain rare [1,2]. Hence, the per sex act transmission probability of Ebola virus from male convalescent survivors is unlikely to be higher than 0.1%, and might well be below this value. Our sensitivity analysis indicates that the duration of the EVD epidemic is heavily influenced by the period during which convalescent men can transmit sexually, calling for a better understanding of the persistence and duration of infectivity of Ebola virus RNA in convalescent patients.

We extended an accepted modeling framework that has been widely used to describe the epidemic trajectories of EVD outbreaks and epidemics [12–14]. To our knowledge, this is the first study using mathematical modeling to assess the potential impact of sexual transmission of Ebola virus on the epidemic in West Africa. In addition, given the generality of the model, this is also the first model that investigates the impact of including a secondary transmission route from convalescent individuals. This model, then, may also have implications for other pathogens with this kind of secondary transmission route (e.g. some adenoviruses [33]; see [34] for examples across mammal species) including other neglected tropical diseases, such as African sleeping sickness [35], and other hemorrhagic fevers that display pathologies similar to EVD [9,10].

In the absence of a better understanding of sexual transmission of EVD, mathematical modeling currently remains the only tool to explore its potential impact on the epidemic trajectory. There are several extensions to this work that could be considered, including transmission by other fluids (e.g., vaginal secretions [4,7]) and asymptomatic infection [36,37], considering spatial aspects of sexual transmission [38], and heterogeneity in sexual behavior [3,39]. We assumed the duration of convalescence to be exponentially distributed. Eggo *et al*. [40] fitted a series of unimodal distributions to the data on Ebola virus RNA detection in semen recently reported by Deen *et al*. [1] and found that the convalescence period could be best described by a gamma distribution. Furthermore, Deen *et al*. [1] found that the cycle-threshold values decreased over time, indicating that the Ebola virus load in semen and the viral infectivity might also decrease during the convalescence period. Uncertainty in the data is not limited to sexual transmission; we fitted our model to weekly incidence of confirmed and probable cases in Sierra Leone, but did not take into account potential underreporting as others have done recently [36]. In addition, EVD is known to exhibit superspreading characteristics [41,42], and superspreading events could lead to explosive regrowth of the epidemic after the occurrence of a new case through sexual transmission [41]. Finally, like other negative-sense single-stranded RNA ((-)ssRNA) viruses [43], the species currently circulating in West Africa has been estimated to have high substitution rates [26,44,45]. This rapid evolution detected throughout the current outbreak zone suggests that within- or between-host adaptation of the virus leading to pro-longed persistence in the seminal fluids is possible. However, evolution of sexual transmission becoming the primary route of spread is highly unlikely.

Awareness of the potential for sexual transmission has led to WHO issuing recommendations that ask convalescent men to abstain from sexual activity as much as possible and to use condoms for up to 6 months after the onset of symptoms [28]. Condom use and social awareness of the risks of sexual transmission during convalescence should have an impact on the per sex act transmission probability (η) and the frequency of sex acts (*q*), respectively. Our results show that condom use should reduce the number of sporadic sexual transmission events during the tail of the epidemic and after discharge of all remaining symptomatic individuals. However, the time during which the public health community must stay vigilant is not reduced because these interventions will not affect the time during which convalescent survivors can shed infectious virus (1/*α*). This is especially poignant since adherence to these recommendations will never be 100%. Thus, our results suggest that the current requirement for declaring a region free from EVD (42 days following either death or a second negative RT-PCR test of the blood from the last known patient), officially declared in Sierra Leone on 7 November 2015 [46], may be premature.

As more data about the convalescent survivors of EVD becomes available, this and future mathematical modeling studies will help to better understand the potential epidemiological consequences of sexual transmission on the EVD epidemic in West Africa. Precise estimates are important for providing convalescent survivors with sound advice that balances protection of the community with the harm that could come from unnecessary stigmatization [47–49].

## Acknowledgements

We thank Benjamin Roche for helpful suggestions on simulation results.

